# LEUTX mediates amplification of DUX4 signaling and aberrant myotube differentiation in facioscapulohumeral muscular dystrophy

**DOI:** 10.64898/2026.10.27.762079

**Authors:** Xiangduo Kong, Sung Jun Beck, Nam Viet Nguyen, Elisabeth Rebboah, Ali Mortazavi, Kyoko Yokomori

**Author notes:** Equal contributions (Xiangduo Kong, Sung Jun Beck).

## Abstract

Aberrant reactivation of an embryonic DUX4 transcription factor gene encoded in the D4Z4 macrosatellite repeats on chromosome 4q is considered the root cause of facioscapulohumeral muscular dystrophy (FSHD). While DUX4 target gene expression correlates with the clinical severity, *DUX4* expression itself is very rare. How this limited *DUX4* expression leads to definitive target gene activation and how these downstream genes contribute to FSHD pathology remain elusive. We previously found the evidence for *LEUTX* and *ZSCAN4* representing two distinct DUX4 target subpathways. Here we present functional analyses in control and FSHD skeletal myocytes identifying LEUTX, but not ZSCAN4, as a major amplifier of DUX4 target gene expression. Furthermore, LEUTX uniquely disrupts myotube differentiation by impairing muscle gene induction and promoting persistent mitotic gene expression. Forced LEUTX expression causes aberrant DNA replication in myotube nuclei. A subset of misregulated genes is bound by LEUTX, strongly suggesting that they are direct targets of LEUTX. Notably, LEUTX requires the presence of DUX4 to gain access to DUX4 target gene promoters for their activation. Taken together, our results establish the critical role of LEUTX in DUX4 signal amplification and defective myotube differentiation, revealing distinct contributions of DUX4 target gene pathways in FSHD pathogenesis.

## Introduction

Facioscapulohumeral muscular dystrophy (FSHD) is a common muscular dystrophy with prevalence of 1 in 8,333 to 20,000 individuals (Deenen et al. 2014). FSHD patients experience weakness in muscles of the face, shoulder girdles and upper arms, which often worsens and spreads to other muscles with age (Statland and Tawil 2016; Wang and Tawil 2021). There are two types of FSHD: FSHD1 (MIM 158900) and FSHD2 (MIM 158901) with similar clinical phenotypes with distinct genetics (de Greef et al. 2010). FSHD1 is characterized by monoallelic contraction of D4Z4 macrosatellite repeats (3.3 kb) at the subtelomeric region of chromosome 4 (4qter), which represents the majority of FSHD patients (> 95 %) (van der Maarel and Frants 2005; van der Maarel et al. 2011). While healthy individuals possess 11 – 150 copies of D4Z4 repeats, FSHD1 patients exhibit 1 – 10 copies. FSHD2 is a rare type of FSHD (≤ 5%) with relatively short D4Z4 repeats (8 – 20 D4Z4 repeats) (Lemmers et al. 2012; Wang and Tawil 2021). The majority of FSHD2 patients exhibit mutations in *SMCHD1*, which is also found to be a disease modifier gene for FSHD1 (Lemmers et al. 2012; Sacconi et al. 2013; Larsen et al. 2015). Mutations of *DNMT3B* and *LRIF1* were also found in FSHD2 cases (van den Boogaard et al. 2016; Hamanaka et al. 2020). Because D4Z4 repeats are primate-specific, the generation of animal models has not been straightforward, hampering a deeper mechanistic understanding of the disease. To this end, we recently developed FSHD mutant myoblast lines carrying CRISPR-engineered *D4Z4* deletion and/or *SMCHD1* mutation, which closely recapitulate the patient cell phenotype and can be compared to the isogenic parental control cells (Kong et al. 2024).

A common feature of FSHD1 and FSHD2 is aberrant expression of the double homeobox 4 (DUX4) transcription factor (Lemmers et al. 2010; Geng et al. 2012; Statland and Tawil 2016; Schätzl et al. 2021). The *DUX4* gene is embedded in each D4Z4 repeat (Gabriëls et al. 1999; Snider et al. 2009). DUX4 is normally expressed transiently during zygotic genome activation (ZGA), the first wave of gene expression during early embryogenesis, and is silenced in somatic tissues with the exception of testis (De Iaco et al. 2017; Whiddon et al. 2017; Karpukhina et al. 2021). We and others demonstrated that loss of heterochromatin (DNA methylation and histone H3 lysine 9 trimethylation (H3K9me3)) at *D4Z4* repeats contributes to abnormal reactivation of *DUX4* in FSHD (van Overveld et al. 2003; Zeng et al. 2014; Kong et al. 2024). Although different isoforms can be expressed from the *DUX4* locus, the full length DUX4 (DUX4fl) containing both transactivation and DNA binding domains is linked to FSHD pathogenesis (Snider et al. 2010; Geng et al. 2012). The “permissive” 4qA haplotype that provides canonical poly-adenylation (poly-A) signal downstream of the distal D4Z4 repeat appears to be required for the disease development, further supporting the significance of functional DUX4 mRNA and protein expression (Dixit et al. 2007; Snider et al. 2009; Lemmers et al. 2010). Upregulation of DUX4 target genes is detected in patient muscles correlating with the clinical severity (Wang et al. 2019; Wong et al. 2020), strongly suggesting that abnormal DUX4 gene network activation is a central driver of FSHD pathogenesis. Since the forced overexpression of DUX4 is highly cytotoxic in myocytes *in vitro* and *in vivo* (Bosnakovski et al. 2008; Vanderplanck et al. 2011; Wallace et al. 2011), DUX4-induced toxicity was thought to be the major driver of the disease (Le Gall et al. 2020; Himeda and Jones 2022). However, the endogenous *DUX4* expression is very rare occurring in less than 1% of patient myoblasts and 3-4% of patient myotubes *in vitro* (Snider et al. 2010; Jones et al. 2012; Tassin et al. 2013; Rickard et al. 2015; Chau et al. 2021). Through bulk and single-nucleus transcriptomic analyses of both primary and immortalized FSHD patient myocytes in vitro, we found little evidence of cell death or apoptotic gene expression in FSHD late myotubes despite higher DUX4 target gene expression than in early myotubes, arguing against a simple model of DUX4 cytotoxicity driving the disease (Jiang et al. 2020; Chau et al. 2021; Kong et al. 2024). Therefore, how the rare DUX4 expression results in definitive disease phenotype was unclear.

DUX4 target genes are more readily detectable in patient muscle cells and have been used as biomarkers of the disease (Lemmers et al. 2010; Geng et al. 2012; Jones et al. 2012; Jiang et al. 2020; Wong et al. 2020; Banerji and Zammit 2021; Chau et al. 2021). We previously found that DUX4 target transcription factor LEUTX contributes to upregulation of DUX4 target genes (Jiang et al. 2020; Chau et al. 2021). We further obtained the evidence that LEUTX and DUX4 target histone variants H3.X/Y amplify each other and are involved in coherent feedforward amplification of the DUX4 gene network activation in FSHD myotubes (Kong et al. 2024). H3.X/Y was shown to be incorporated into transcriptionally active gene loci and maintain their open state, contributing to increased inducibility of DUX4 target genes (Resnick et al. 2019). However, the mechanistic detail of LEUTX action was unclear. Furthermore, our recent MERFISH spatial-transcriptomic analyses suggested that DUX4 targets are unequally expressed and that there appear to be at least two DUX4 target subpathways represented by the *LEUTX* and *ZSCAN4* groups in FSHD myotubes (Chen et al. 2024). Thus, dissecting the DUX4 target gene network and delineating specific contributions of individual DUX4 targets and subpathways are critical to understand FSHD pathogenesis. In particular, the *LEUTX* gene is missing in rodents and is not possible to study in the mouse model. Here, we examined the effect of LEUTX and ZSCAN4 expression using FSHD patient and control myocytes, as well as isogenic mutant myocytes carrying the *D4Z4* contraction and *SMCHD1* mutation associated with severe FSHD1. Our results distinguish the functions of these two genes and identify LEUTX as a DUX4-dependent DUX4 target gene amplifier that also interferes with normal muscle differentiation, providing a mechanistic link between DUX4 network activation and FSHD muscle pathology.

## Results

### Distinct contributions of LEUTX and ZSCAN4 in DUX4 target gene expression

Using lentiviral shRNAs, we depleted LEUTX or ZSCAN4 in immortalized FSHD patient and mutant cell lines at the early myotube stage (Fig. 1A). The FSHD2 patient cell line used (B2) carries both the *SMCHD1* mutation and a modest contraction of D4Z4 repeats, with two 4qA alleles containing 15 and 19 D4Z4 repeats. This cell line has been used as a representative of FSHD cells as it expresses relatively high FSHD-induced genes (most of which are DUX4 target genes) as previously characterized (Jiang et al. 2020; Chau et al. 2021; Kong et al. 2024; Kong et al. 2025). The mutant cell line was derived from the healthy control line (B0) and is therefore isogenic. It carries the *D4Z4* deletion and *SMCHD1* mutation and expresses FSHD-induced and DUX4 target genes, providing a complementary disease model (Kong et al. 2024). We performed bulk RNA-seq and hierarchical clustering of gene expression changes using fold-change values across 61 FSHD/DUX4-induced genes (Jiang et al. 2020), most of which have confirmed DUX4 protein binding to the promoter regions (Yao et al. 2014; Kong et al. 2024) (Fig. 1B). We found that LEUTX depletion results in global downregulation of these genes in FSHD (patient) myotubes similar to mutant myotubes as previously shown (Supplemental Fig. S1A) (Kong et al. 2024). In contrast, ZSCAN4 depletion downregulated only a subset of FSHD-induced genes, and unexpectedly, upregulated a few other target genes, such as *H3.X*, *DUXA* and *PRAMEF13*, displaying a phenotype distinct from LEUTX depletion. RT-qPCR reproduced these results, and the effects of LEUTX depletion were confirmed in additional FSHD cell lines (Supplemental Fig. S1B and C).

**Figure 1.**
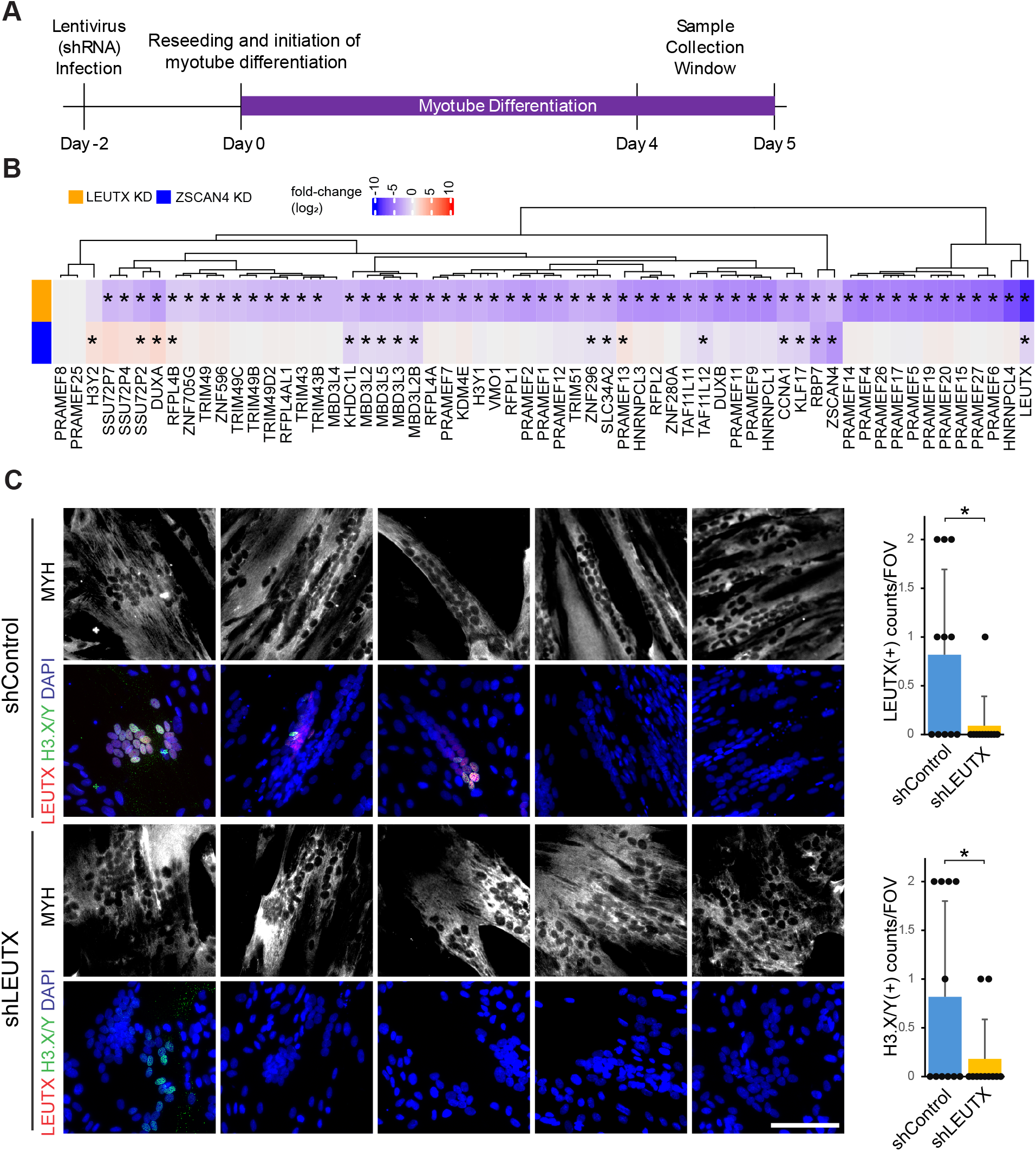
LEUTX depletion results in the downregulation of more DUX4 targets compared to ZSCAN4 depletion. **A.** The schematic diagram for the depletion experiments is shown. **B.** A heatmap of fold-change values obtained from the bulk RNA-seq experiments between the LEUTX depletion and the ZSCAN4 depletion in the FSHD2 (B2) myotubes compared to the control shRNA-treated cells. The heatmap shows 61 pre-selected DUX4 targets where asterisks indicate statistically significant (q-value < 0.1) fold-change values. **C.** 20X immunofluorescence representative images of the FSHD2 (B2) myotubes treated with the control shRNA (shControl) and the LEUTX shRNA (shLEUTX) where MYH, H3.X/Y and LEUTX are presented in gray, green and red colors. Scale bar = 100 μm. Five different fields of view are shown as representative images. Upper right panel: Quantification of LEUTX+ myotube counts per field of view between the control shRNA (shControl) and the LEUTX shRNA (shLEUTX) is shown. Lower right panel: The same quantification is shown for H3.X/Y+ myotube counts. Zero-Inflated Poisson model was used. * p ≤ 0.05. n=11.

The histone variants H3.X/Y are shown to be upregulated by DUX4 and incorporated into actively transcribed gene regions to promote their expression (Resnick et al. 2019). We have previously shown that *H3.X/Y* and *LEUTX* reciprocally enhance their expression at the RNA level (Kong et al. 2024). LEUTX and H3.X/Y proteins colocalize well in the myotube nuclei (Supplemental Fig. S1D), and depletion of LEUTX by shRNA led to reduction of both proteins (Fig. 1C). Taken together, the results identify LEUTX as a major amplifier of DUX4 target gene activation.

### LEUTX blocks downregulation of mitosis-related gene expression in FSHD early myotubes

Including DUX4 target genes, we found 302 genes that are commonly downregulated (Log_2_(fold change) < -1 and q-value < 0.1) by LEUTX depletion in both patient and mutant myotubes (Fig. 2A, top) (Supplemental Table S1). In these 302 genes, we found the top GO terms are biological processes associated with mitosis and cell proliferation. We selected the top 10 significantly enriched terms and compiled the 75 unique genes that comprised these terms, hereafter referred to as “mitosis-related” genes (Fig. 2B, top) (Supplemental Table S2). As expected, we found that only LEUTX depletion downregulated these mitosis-related genes, whereas ZSCAN4 depletion had no effect (Fig. 2C, yellow circles). Representative genes include *INCENP, CDC25C, NUSAP1, FBXO5 and CENPF*, as indicated (Fig. 2C, yellow circles). Upon myoblast fusion and differentiation into myotubes, mitotic and cell cycle genes are expected to be downregulated (Moran et al. 2002; Shen et al. 2003). Indeed, these mitosis-related genes were effectively downregulated upon myotube differentiation in healthy control cell lines (Fig. 2D, left, compare myoblasts (MB) and myotubes (MT)). In contrast, downregulation of these genes is weaker in both patient (FSHD1 and FSHD2) and mutant myotubes (Fig. 2D, left). These results show that LEUTX, but not ZSCAN4, sustains mitosis-related gene expression during myotube differentiation in FSHD cells.

**Figure 2.**
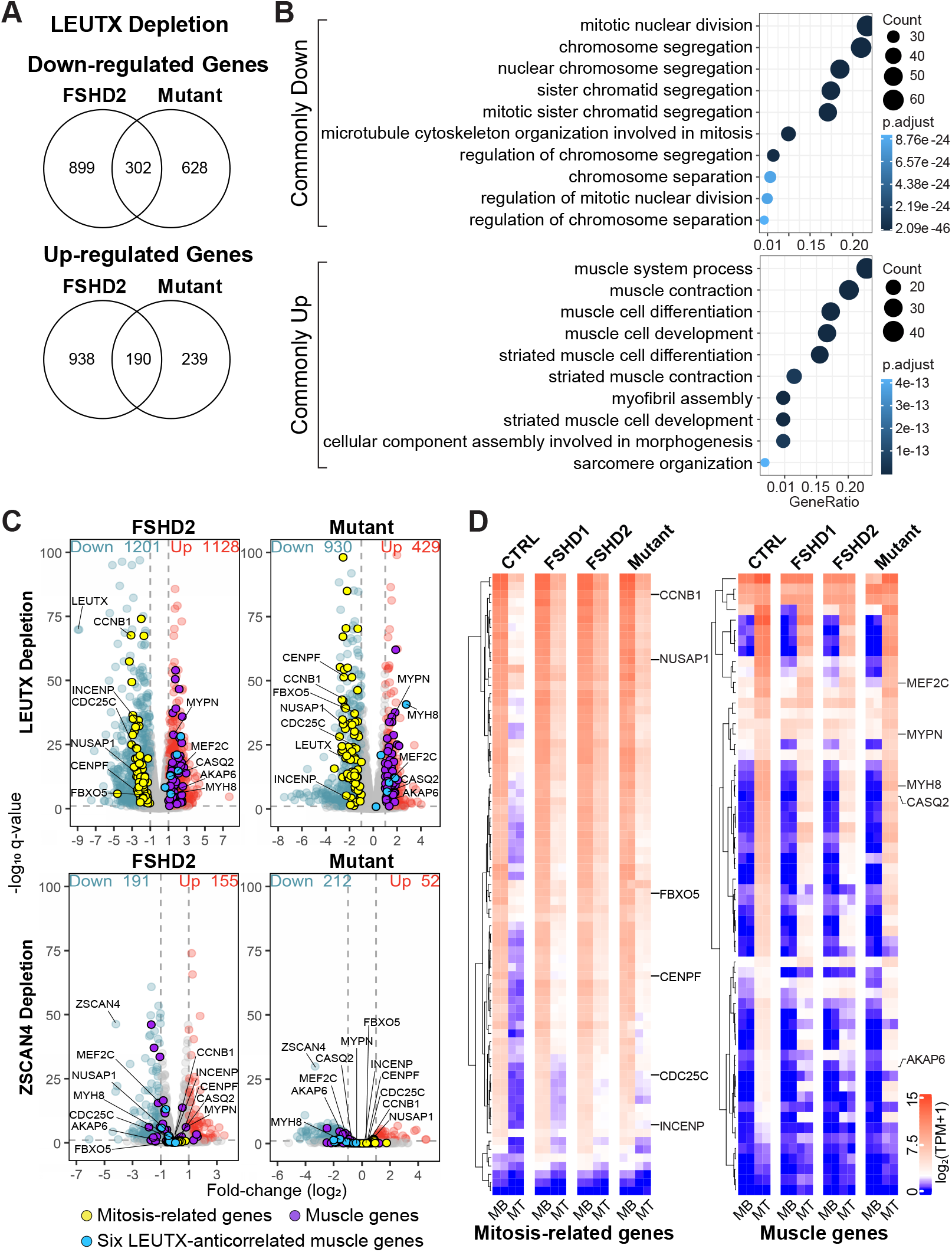
LEUTX depletion, but not ZSCAN4 depletion, decreases mitosis-related genes and increases muscle gene expression in the FSHD context. **A.** Venn diagrams indicate the number of downregulated and upregulated DEGs by LEUTX depletion in the FSHD2 and the mutant cell lines. **B.** Enriched GO terms from the commonly downregulated (top panel) and commonly upregulated (bottom panel) DEGs are shown by dot plots. **C.** Volcano plots indicate genes associated with mitosis (yellow dots) and muscle contraction and development (purple dots). Text labels indicate representative mitosis-related genes such as *CDC25C*, *NUSAP1*, and *INCENP*, and muscle genes such as *CASQ2*, *MEF2C*, and *MYH8*. Light blue dots indicate six muscle genes that previously showed negative correlations with *LEUTX* expression in MERFISH analysis (Chen et al. 2024). **D.** TPM values of the mitosis-related genes (left) and muscle genes (right) identified in (**C**) in previously published RNA-seq data of control, representative FSHD cells (FSHD1 and FSHD2) and double mutant cells at myoblast and early myotube stages (Kong et al. 2024) are shown as indicated. Representative genes are labeled.

### LEUTX interferes with upregulation of muscle-related genes in early FSHD myotubes

Using the same approach, we found 190 genes are commonly upregulated (Log_2_(fold change) > 1 and q-value < 0.1) by LEUTX depletion in both patient and mutant myotubes (Fig. 2A, bottom) (Supplemental Table S3). GO term analyses of commonly upregulated genes showed strong enrichment for muscle contraction (Fig. 2B, bottom) (Supplemental Table S4). We selected the top 12 most significantly enriched GO terms and compiled the 60 unique genes that comprised these terms, hereafter referred to as “muscle-related” genes. Those genes are not upregulated by ZSCAN4 depletion, and some were downregulated by ZSCAN4 depletion in FSHD cells, further highlighting the distinct transcriptomic phenotypes induced by depletion of LEUTX and ZSCAN4 (Fig. 2C, purple circles).

Our previous MERFISH analyses revealed strong negative expression correlations between the endogenous *LEUTX* expression and six muscle genes, *MYOM3, MEF2C, TRIM72, CASQ2, MYH8* and *LAMA5*, in FSHD patient and mutant myotubes (Chen et al. 2024). We found that these genes are indeed upregulated by LEUTX depletion (Fig. 2C, blue circles). Impaired upregulation of muscle-related genes in FSHD and DUX4-inducible cells has been reported in multiple studies (Winokur et al. 2003; Tsumagari et al. 2011; Geng et al. 2012; Jagannathan et al. 2016; Kong et al. 2024). Consistent with these reports, muscle-related genes, such as *MEF2C, MYPN, MYH8*, *CASQ2* and *AKAP6,* were clearly induced in control cells at the early myotube stage, but showed variable and impaired upregulation in patient and mutant myotubes (Fig. 2D, right). Taken together, our results reveal that LEUTX interferes with mitotic gene silencing and muscle gene induction, contributing to the FSHD transcriptomic phenotype.

### LEUTX overexpression interferes with myotube differentiation without cytotoxicity

DUX4 overexpression was shown to interfere with myoblast differentiation and lead to cell death in vitro (Bosnakovski et al. 2008; Vanderplanck et al. 2011; Jagannathan et al. 2016). We generated stable healthy control and FSHD cell lines with DOX-inducible FLAG-tagged LEUTX (Fig. 3A). We also overexpressed FLAG-tagged ZSCAN4 as comparison. Protein induction was confirmed by Western blot (Fig. 3B). While comparable LEUTX and ZSCAN4 proteins are detectable only in the presence of DOX in control cells, additional faster migrating bands observed without DOX in FSHD cells likely correspond to the endogenous proteins (Fig. 3B). In a differentiation medium, overexpression of LEUTX for 48 hours effectively blocked myotube differentiation in both control and FSHD samples whereas no significant effect of ZSCAN4 expression was observed (Fig. 3C, Supplemental Fig. S2). The same phenotype in healthy control cells demonstrates that LEUTX expression alone, in the absence of DUX4, is sufficient to impair differentiation. Because myogenic differentiation normally requires cell cycle exit, we measured 5-ethynyl-2’-deoxyuridine (EdU) incorporation as a marker for ongoing DNA synthesis. Following 24 hours of doxycycline (DOX) induction started on Day 3 of differentiation, LEUTX expression was not sufficient to interfere with myotube fusion, but significantly increased EdU-positive cells compared to no DOX treatment, suggesting that LEUTX stimulates DNA replication during early differentiation (Fig. 3D).

**Figure 3.**
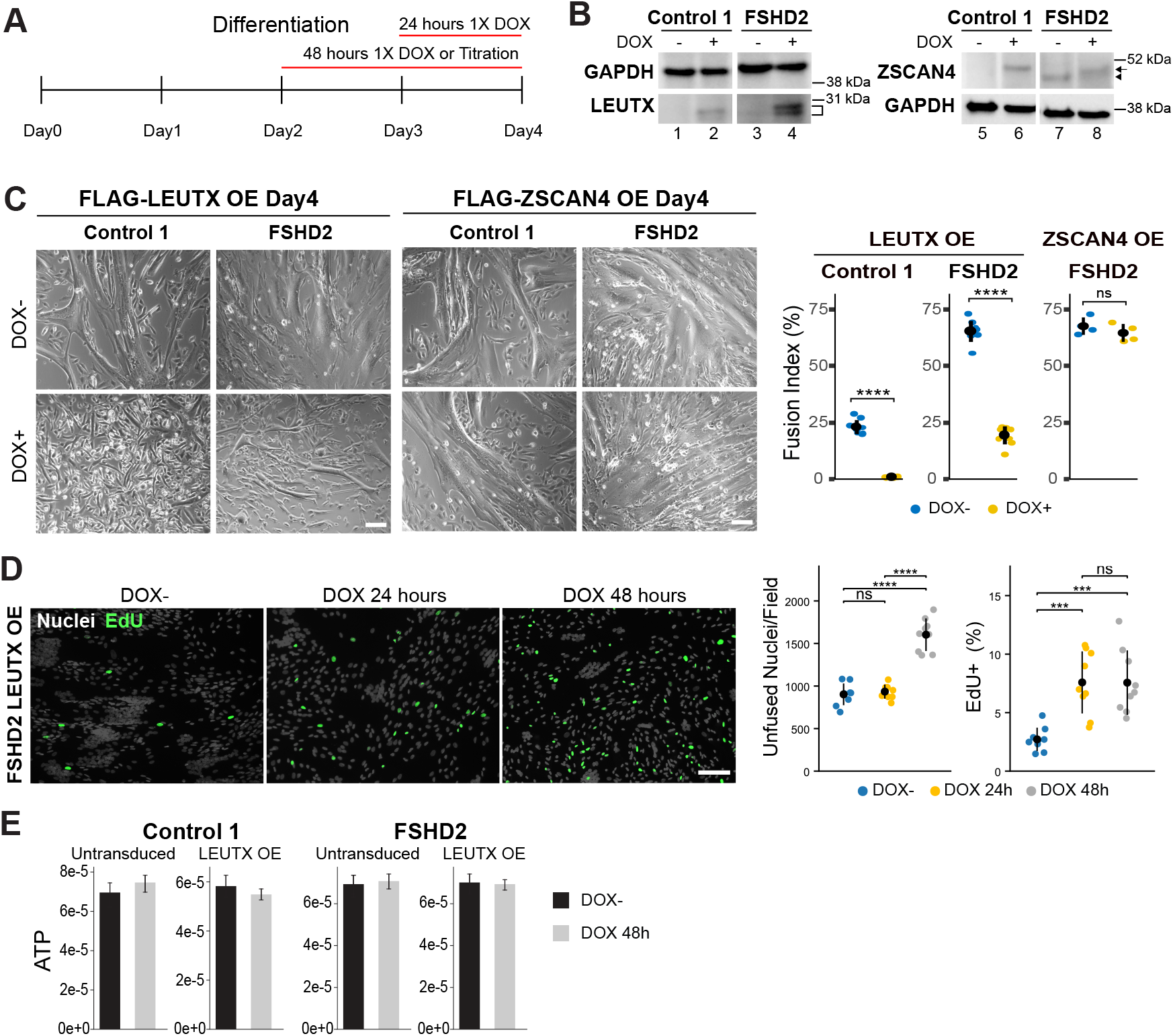
LEUTX overexpression interferes with myotube differentiation without cytotoxicity. **A.** The schematic diagram indicates the timeline of LEUTX overexpression conditions. **B.** Western blot images of LEUTX or ZSCAN4 protein bands are shown where GAPDH was used as the loading control. “–” indicates no DOX condition and “+” indicates 48 hours DOX treatment. **C.** 10X phase contrast representative images of the healthy control 1 (B0) and the FSHD2 myotube differentiation in Day 4 with and without 48 h DOX treatment for LEUTX or ZSCAN4 overexpression are shown. Scale bar 100 µm. Fusion index % of the healthy control 1 and the FSHD2 myotube between with and without DOX treatment is shown in the right panel. Student’s t-test. **** p ≤ 0.0001. n=9 (LEUTX overexpression). ns not significant. n=4 (ZSCAN4 overexpression). **D.** Representative immunofluorescence images of the FSHD2 are shown, indicating nuclei (counter stain by DAPI) in gray and EdU signals in green. Scale bar = 100 μm. Right panel: Quantification of DAPI stained nuclei and EdU-positive signals under control (DOX−), 24 h DOX (1×), and 48 h DOX (1×) treatment conditions. Student’s t-test. *** p ≤ 0.001 **** p ≤ 0.0001. n=9. **E.** The effect of 1X DOX concentration was tested for cell viability, measuring ATP level between the untransduced parental cells and the transduced stable cell lines. 1X DOX concentration was tested on the healthy control 1 (B0) (n=4) and the FSHD2 (B2) (n=6). Student’s t-test. All comparisons resulted in no statistical significance.

We measured the ATP level as a proxy of cell viability with and without LEUTX overexpression in control and FSHD cell lines. Unlike the toxicity of DUX4 overexpression as previously reported (Bosnakovski et al. 2017), LEUTX expression had no effect on cell viability even after 48-hour induction when myotube fusion is significantly blocked (Fig. 3E). Taken together, the results indicate that forced expression of LEUTX in myoblasts interferes with cell cycle exit and myotube fusion without causing cytotoxicity, suggesting that cytotoxicity induced by DUX4 in myoblasts is independent of the LEUTX pathway.

We previously reported that the endogenous *LEUTX* is one of the “late” genes that are activated by DUX4 almost exclusively in differentiated myotubes rather than in myoblasts (Kong et al. 2024). Thus, we analyzed the overexpression of the recombinant LEUTX using transient transduction in differentiated early control 2 (D0) myotubes (Fig. 4A). Interestingly, we found that some of the LEUTX-positive nuclei in DOX+ myotubes coincide with the proliferation marker Ki67 and EdU incorporation whereas no Ki67- and EdU-positive nuclei are found in DOX-myotubes at this stage (Fig. 4B). The results reveal that LEUTX expression in myotubes can trigger DNA replication, indicative of aberrant re-entry into the cell cycle. Taken together, the results indicate that LEUTX expression interferes with proper cell cycle exit and differentiation of myotubes.

**Figure 4.**
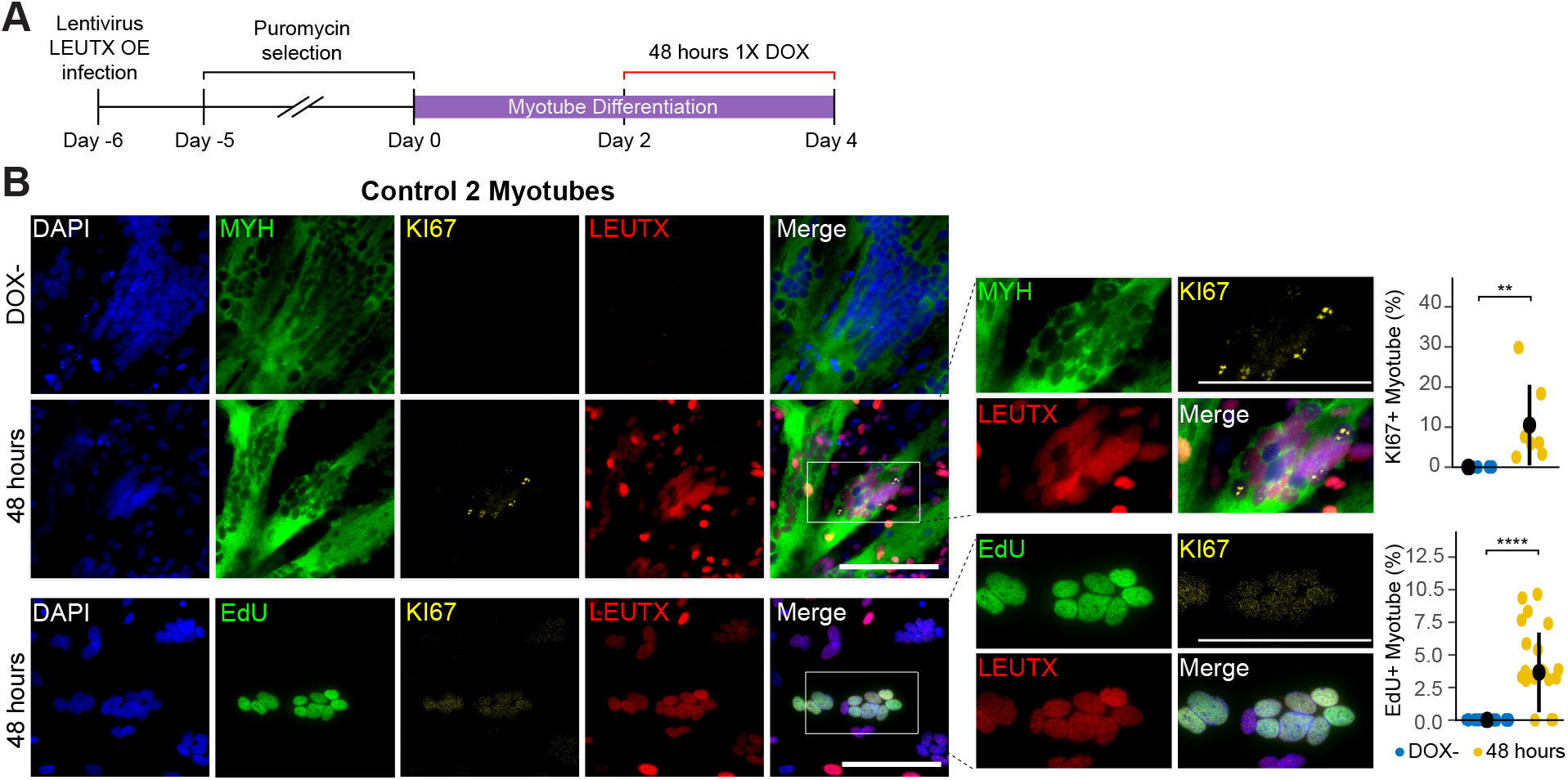
LEUTX overexpression promotes DNA replication and proliferation state in the myotube differentiation environment. **A.** The schematic diagram for the transient LEUTX overexpression experiments. **B.** Representative immunofluorescence images of the healthy control 2 (D0) are shown, indicating nuclei (counter stain by DAPI) in blue, MYH in green, EdU in green, KI67 in yellow, and LEUTX in red. Scale bar = 100 μm. The middle panel shows magnified views of the boxed regions from the corresponding merged images in the left panel. The right panel: Quantification of the KI67+ myotube percentage (%) and EdU+ myotube percentage is shown. Wilcoxon test. ** p ≤ 0.01 n=7 (for the KI67 quantification). **** p ≤ 0.0001 n=25 (for the EdU quantification).

### Genome-wide mapping defines LEUTX binding sites in human myocytes

The aberrant myotube differentiation by LEUTX overexpression agrees with the transcriptomic phenotype of FSHD patient and mutant cells; muscle genes are not properly upregulated and cell cycle/mitosis genes remain high, which was reversed by LEUTX depletion (Fig. 2C and D). This raises the possibility that LEUTX may directly activate mitosis-related genes and repress muscle-related genes. To address which genes may be directly controlled by LEUTX, we mapped LEUTX occupancy by CUT&Tag at day 4 of differentiation in two complementary systems: control myocytes overexpressing recombinant LEUTX (control LEUTX OE) as well as in FSHD1 (C1) cells with high expression of DUX4 targets, including the endogenous LEUTX (FSHD enLEUTX) (Chau et al. 2021) (Fig. 5). Using a polyclonal LEUTX antibody, we identified 18,327 reproducible binding sites in control LEUTX OE and 12,520 binding sites in FSHD enLEUTX (Fig. 5A; Supplemental Fig. S3A). To confirm the antibody specificity, we also used a monoclonal antibody specific for LEUTX and obtained weaker but concordant peaks for FSHD enLEUTX, validating the results with the polyclonal antibody (Supplemental Fig. S3B and C). Notably, there are only 2,543 overlapping peaks between control LEUTX OE and FSHD enLEUTX, suggesting the context-dependent binding of LEUTX in the FSHD background (Fig. 5A; Supplemental Fig. S3C).

**Figure 5.**
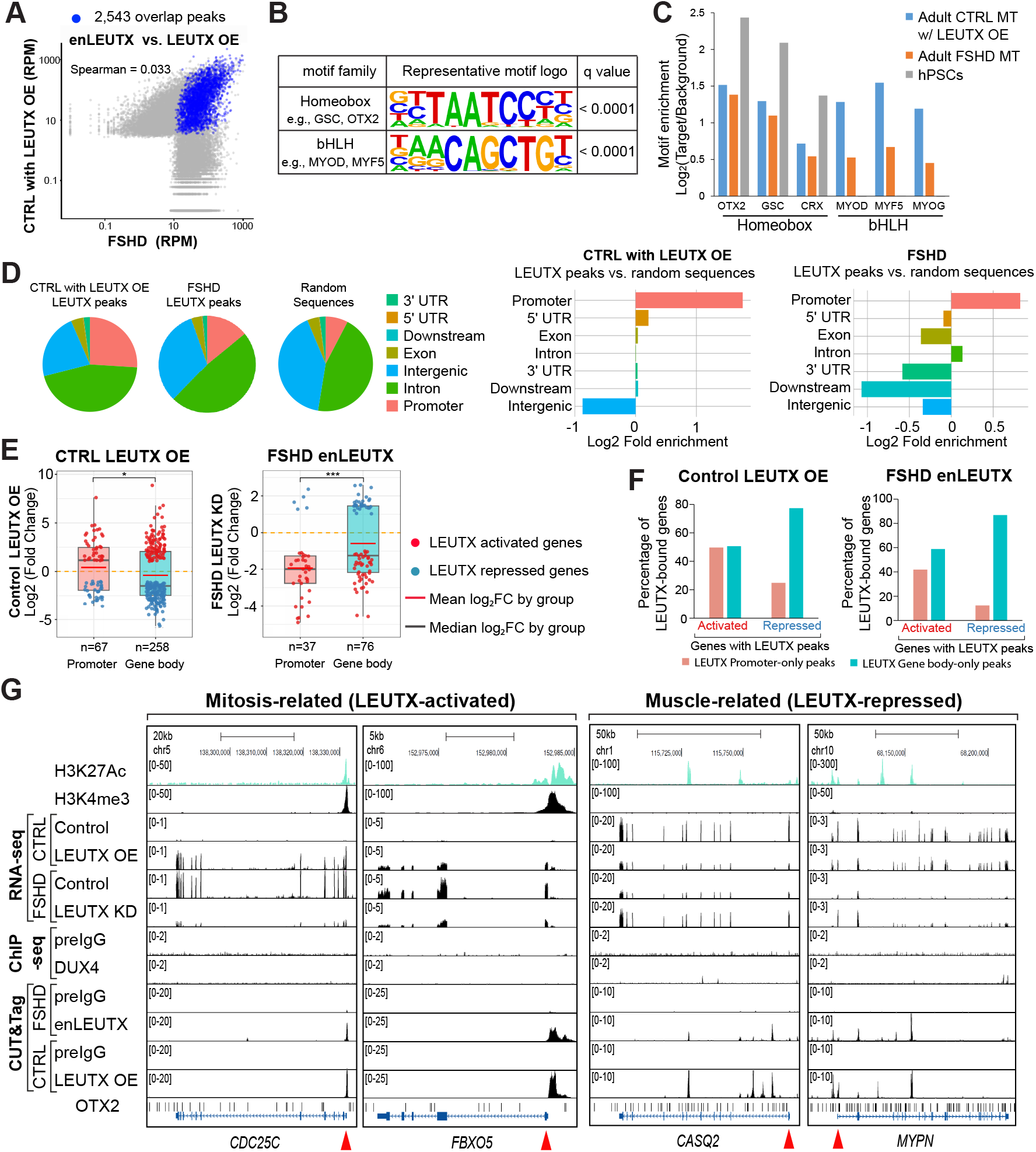
LEUTX-binding sites identified LEUTX functions as both a transcriptional activator and repressor. **A.** Scatter plot showing LEUTX CUT&Tag peak counts (RPM), and the correlation of peak intensities between DOX-inducible LEUTX control myocytes and FSHD myocytes. Each point represents an individual LEUTX peak, with blue dots indicating peaks shared between the two conditions. **B.** Known motif enrichment analysis of LEUTX-binding sites using HOMER shows enrichment of homeobox and bHLH motif families. Representative motif logos for each motif family (OTX2 and MYF5). **C.** Motif enrichment analysis of LEUTX CUT&Tag/ChIP peaks in adult control myotubes with LEUTX OE, adult FSHD myotubes, and human pluripotent stem cells (hPSCs). Enrichment is shown as log_2_ transformed target-to-background ratios for representative homeobox (OTX2, GSC, and CRX) and bHLH (MYOD, MYF5, and MYOG) motifs. **D.** Genomic distribution of LEUTX peaks across annotated genomic regions. A control dataset consisting of randomly selected genomic regions matched for peak number and length was generated 1,000 times, and the averaged distribution was used for comparison. Promoter regions are defined as 2,500 bp upstream and 500 bp downstream of the transcription start site (TSS). Downstream regions are defined as 500 bp upstream of the transcription termination site (TTS) and 2,500 bp downstream of the TTS. The 5’ and 3’ untranslated regions (UTRs) were defined based on UCSC genome annotations, excluding the 500 bp immediately adjacent to the TSS or TTS, respectively. When a peak overlapped multiple genomic annotations, it was assigned to a single category based on the following priority order: promoter, 5’ UTR, intron, exon, 3’ UTR, downstream, and intergenic. Right panel: Enrichment of LEUTX peaks across genomic regions relative to randomly sampled genomic sequences. **E.** Gene expression changes associated with LEUTX-bound regions located in promoter or gene body regions. Boxplots show the distribution of log₂ fold changes in gene expression for genes associated with LEUTX peaks located in promoter or gene body regions in control cells with LEUTX OE (left) and FSHD cells following LEUTX KD (right), as indicated in the figure. Each dot represents an individual gene associated with a LEUTX peak. Sample sizes (n) represent the number of genes associated with LEUTX peaks in each genomic region. Statistical significance between promoter- and gene body-associated genes was assessed using an unpaired two-tailed t-test (*p < 0.05, ***p < 0.001). **F.** Distribution of LEUTX CUT&Tag peaks across promoter and gene body regions in LEUTX-activated and LEUTX-repressed genes, as shown in (**E)**. **G.** UCSC Genome Browser snapshots showing LEUTX binding and associated chromatin features at representative mitosis-related genes (left 2 panels) and muscle-related genes (right 2 panels). Tracks include H3K27Ac (ENCODE), H3K4me3 (our control cells), RNA-seq signals from control cells with and without LEUTX OE as well as FSHD myotubes with control shRNA or LEUTX shRNA (LEUTX KD) treatment, control pre-IgG and DUX4 ChIP-seq signals (Geng et al. 2012), as well as CUT&Tag signals using polyclonal antibody specific for LEUTX for FSHD enLEUTX and control LEUTX OE with corresponding pre-IgG, as indicated.

LEUTX was previously shown to be a transcriptional activator that binds to the homeobox OTX2 motif (“TAATCC”) in pluripotent stem cells (PSCs) (Jouhilahti et al. 2016; Katayama et al. 2018; Gawriyski et al. 2023). We therefore asked whether LEUTX shows a similar binding preference in myocytes. Interestingly, the motif analysis revealed almost equally significant enrichment for the homeobox motifs (e.g., OTX2) and the bHLH motifs (e.g., MYF5/MYOD/MYOG) with control LEUTX OE, and less but also significant enrichment of bHLH for FSHD enLEUTX (Fig. 5B and C). In contrast, LEUTX ChIP-seq in PSCs (Gawriyski et al. 2023) showed homeobox motif enrichment but does not yield any enrichment for MYF5, MYOD and MYOG motifs (Fig. 5C). These results indicate that LEUTX retains its known homeobox motif preference while acquiring a muscle-associated binding pattern in skeletal myocytes.

### LEUTX functions as both a transcriptional activator and repressor

To determine how LEUTX binding relates to its effects on gene expression, we integrated the CUT&Tag chromatin occupancy data with RNA-seq from control cells with and without LEUTX OE and FSHD cells with and without LEUTX knockdown. In control cells, genes increased by LEUTX OE were classified as LEUTX-activated while genes decreased by LEUTX OE were classified as LEUTX-repressed. In FSHD cells, direction was inferred by LEUTX depletion, where genes that decreased following LEUTX depletion were classified as LEUTX-activated and genes that increased were classified as LEUTX-repressed. It should be noted that there was no significant upregulation of embryonic genes by LEUTX OE in the context of myotubes (Supplemental Fig. S4) (Yan et al. 2013; Zheng et al. 2024).

Genome-wide analysis shows that LEUTX binding peaks for both control LEUTX OE and FSHD enLEUTX are highly enriched at the promoters relative to matched random genomic regions (Fig. 5D). Genes with LEUTX binding at promoter regions are more frequently activated than repressed in both control LEUTX OE and FSHD enLEUTX cells (Fig. 5E). Downstream gene body binding was observed among both LEUTX-activated and repressed targets (Fig. 5E). Notably, LEUTX-repressed genes showed strong enrichment for peaks in the gene body in both control LEUTX OE and FSHD enLEUTX (Fig. 5F). Together, the results associate LEUTX promoter binding primarily with gene activation and identify gene body binding as a common feature of LEUTX-repressed targets.

We next asked which differentially expressed genes following LEUTX depletion in FSHD myotubes are directly bound by LEUTX. Of the 302 LEUTX-activated genes (downregulated by LEUTX depletion in FSHD cells), 42% contain LEUTX peaks in control LEUTX OE cells, compared to 33% in FSHD with enLEUTX cells (Supplemental Figure S5). Of the 190 LEUTX-repressed genes (activated by LEUTX depletion in FSHD cells), 49% contained LEUTX peaks in control LEUTX OE, compared with 24% in FSHD enLEUTX (Supplemental Figure S5). These include 44% of mitosis-related (LEUTX-activated) and 63% of muscle-related genes (LEUTX-repressed) in LEUTX OE control, compared with 24% and 27%, respectively, in FSHD (enLEUTX) cells (Supplemental Figure S5). The lower proportion of LEUTX-bound genes in FSHD cells may reflect lower amount of the endogenous LEUTX expression or differences in cofactor availability and the epigenetic environment. Together, these results support direct regulation of subsets of mitosis-related and muscle-related genes by LEUTX.

Mitosis-related genes, *CDC25C* and *FBXO5,* are induced by LEUTX OE in control cells and repressed by LEUTX depletion in FSHD cells (LEUTX-activated) (Fig. 5G, RNA-seq in the left two panels). We found significant LEUTX binding peaks in the promoter regions of these genes in both control LEUTX OE and FSHD (enLEUTX) cells (Fig. 5G, left 2 panels). Muscle-related genes, *CASQ2* and *MYPN*, are repressed by LEUTX OE in control cells and upregulated by LEUTX depletion in FSHD cells (LEUTX-repressed) (Fig. 5G, RNA-seq in the right 2 panels). These genes indeed show multiple peaks in the downstream gene body regions, often significantly more prominent than promoter peaks when present, strongly suggesting direct repression by LEUTX (Fig. 5G, right 2 panels). LEUTX peaks at the promoters of the LEUTX-activated genes specifically coincide with significant histone H3 lysine 27 acetylation (H3K27Ac) in ENCODE myocyte data whereas discordant and partial overlaps are observed in the gene body regions of the repressed genes (Fig. 5G). Taken together, our results indicate that LEUTX functions as both activator and repressor in human skeletal myotubes, potentially through distinct binding modes and associated mechanisms.

### LEUTX overexpression enhances DUX4 target gene expression through DUX4-induced promoter binding

Consistent with the previous study in mutant cells (Kong et al. 2024) and inversely correlating with the depletion effect in FSHD cells (Fig. 1B), LEUTX overexpression enhanced DUX4-induced gene expression in FSHD cells (Fig. 6A). RT-qPCR results confirmed dose-dependent induction of DUX4 targets in FSHD cells (Supplemental Fig. S6A). We confirmed that LEUTX overexpression does not induce *DUX4* (Kong et al. 2025). The OE effect may be attenuated in FSHD cells because the endogenous LEUTX already stimulates these genes, masking the effect of additional LEUTX expression, compared to more robust and global repression effects observed with LEUTX depletion (Figs. 1B and 6A). LEUTX expression induced *VMO1* in control cells (Fig. 6A and C). *VMO1* is a known LEUTX target gene (Gawriyski et al. 2023) and exhibited strong expression correlation with the endogenous *LEUTX* in our previous spatial transcriptomics study of patient and mutant myocytes (Chen et al. 2024). The results indicate that the majority of DUX4 target gene activation by LEUTX occurs only in FSHD cells.

**Figure 6.**
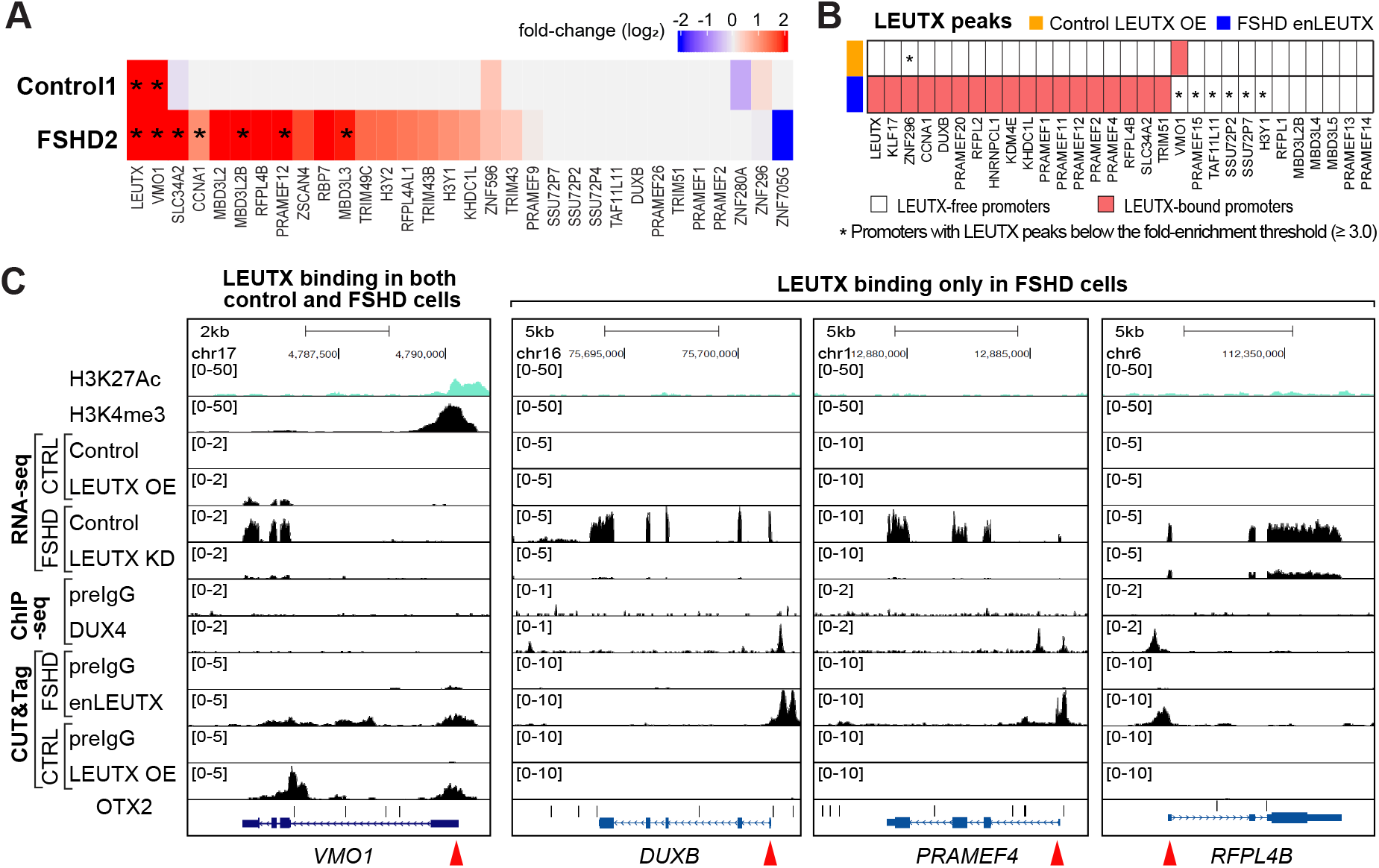
LEUTX enhances DUX4 target gene expression through DUX4-induced promoter binding. **A.** A heatmap of fold-change values obtained from the bulk RNA-seq experiments from the LEUTX overexpression in control and FSHD cell lines. The heatmap shows 33 pre-selected DUX4 targets where asterisks indicate statistically significant (q-value < 0.1) fold-change values. **B.** Schematic representation of LEUTX binding at the promoters of DUX4 target genes in control cells with LEUTX OE and FSHD1 cells with endogenous LEUTX. Red boxes indicate gene promoters with LEUTX CUT&Tag peaks. * indicates regions where the endogenous LEUTX shows peaks that are too weak to make the cut-off (Supplemental Figure S6). **C.** UCSC Genome Browser image of example DUX4-induced genes. Tracks include H3K27Ac (ENCODE), H3K4me3 (our control cells), RNA-seq signals of control cells without and with LEUTX OE as well as control and LEUTX KD in FSHD myotubes, DUX4 ChIP-seq signals (Geng et al. 2012), LEUTX CUT&Tag signals of control LEUTX OE and FSHD enLEUTX with corresponding pre-IgG controls, as indicated. Left panel, DUX4-activated gene that can be bound and induced by LEUTX in control cells without DUX4. Right 3 panels: DUX4 target genes in which LEUTX binds to the promoter and enhances its expression only in FSHD cells.

Consistent with the above results, we found that LEUTX binds to the DUX4 target gene promoters in FSHD, but not in control, myotubes (Fig. 6B). Notably, those promoters, including *VMO1*, that are bound by LEUTX in control cells are marked by H3K27Ac (by ENCODE) as well as H3K4me3 (CUT&Tag done in our control cells), whereas the promoters that LEUTX binds only in FSHD cells are missing these marks in control, suggesting that the latter promoters are normally closed (Figs. 5G and 6C). At these FSHD-specific promoters, LEUTX binding peaks closely coincide with DUX4 binding peaks (Geng et al. 2012) (Fig. 6C). The results suggest that the effect of LEUTX on DUX4 target genes is dictated by its promoter accessibility and that DUX4 binding is prerequisite for LEUTX to gain access to these promoters. This model is consistent with our previous finding that LEUTX depletion hindered DUX4 target gene expression only later in differentiation, whereas initial activation requires DUX4 (Jiang et al. 2020; Chau et al. 2021). Together, these results reveal that DUX4 is not only important for initially activating *LEUTX* expression but is also critical for allowing LEUTX binding to other DUX4 target gene promoters for feed-forward DUX4 target amplification.

## Discussion

While DUX4 reactivation has been linked to FSHD, how the very rare expression of DUX4 leads to widespread muscle pathology remains unresolved. Though recombinant DUX4 overexpression is cytotoxic, our previous bulk and single nucleus transcriptomic analyses argue against overt cytotoxicity of the endogenous DUX4 expression (Jiang et al. 2020; Kong et al. 2024). Rather, our recent analyses began to unravel the critical roles of DUX4 target genes in DUX4 signal amplification and differential contributions to FSHD phenotype (Chen et al. 2024; Kong et al. 2024). Since not all DUX4 target genes are expressed equally, with LEUTX and ZSCAN4 representing different co-expression modules (Chen et al. 2024), we compared their functions in control and FSHD myocytes. This is important as LEUTX is not present and thus its role cannot be assessed in animal models (Yao et al. 2014). Our results reveal the role of LEUTX, not ZSCAN4, as a major amplifier of the DUX4 target gene network and in interfering with myotube differentiation without inducing cytotoxicity (Fig. 7A). The results highlight the unique contributions of LEUTX in FSHD pathology.

**Figure 7.**
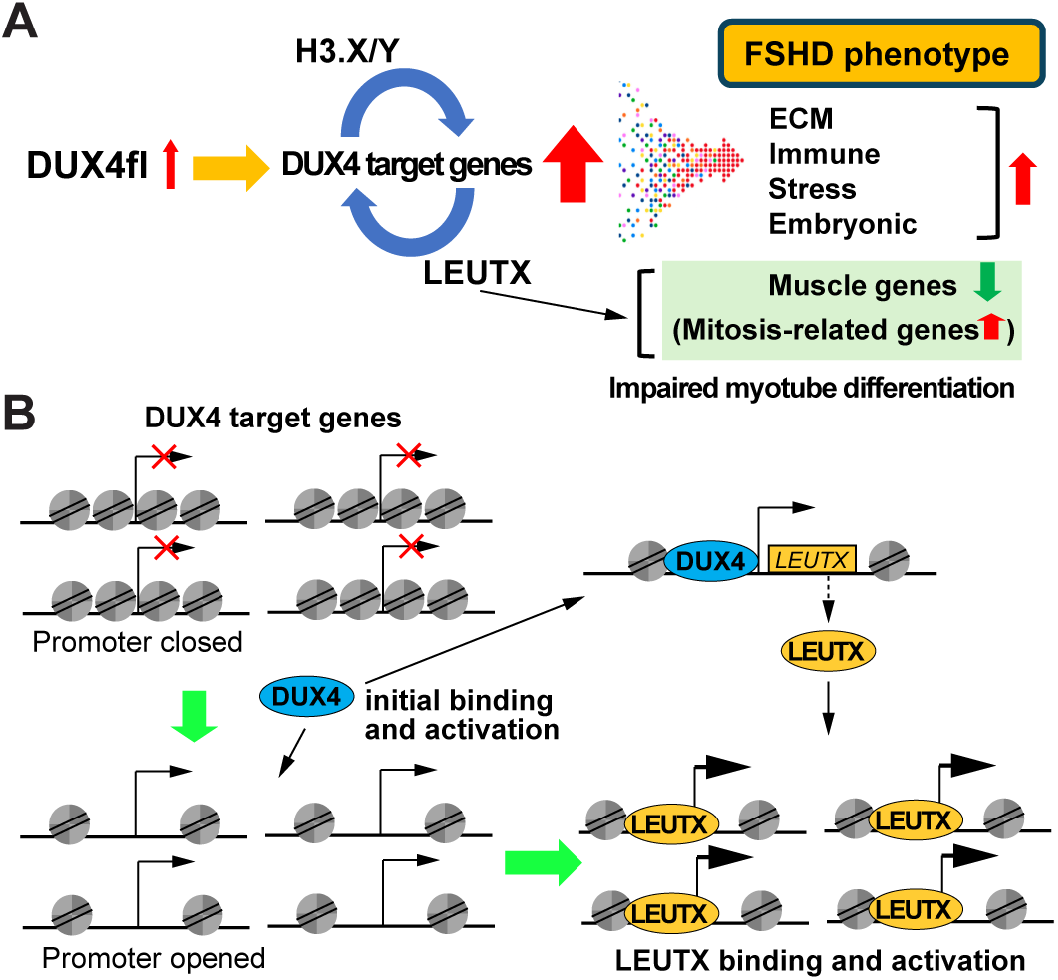
Contributions of LEUTX to FSHD phenotype. **A.** Limited DUX4 activation leads to significant DUX4 target gene expression through feedforward mechanism involving H3.X/Y and LEUTX, resulting in FSHD transcriptomic phenotype. LEUTX specifically interferes with myotube differentiation by muscle-related gene downregulation and mitosis-related gene activation. Downregulation of muscle-related genes has been observed in FSHD patient cells. **B.** A two-step mechanism of DUX4 gene network amplification. DUX4 initially triggers the expression of its target genes, including *LEUTX*, by penetrating and opening the otherwise closed chromatin at the target gene promoters as a disease-specific pioneer factor. LEUTX protein then gains access to and activates DUX4 target genes.

### Differential contributions of DUX4 downstream target genes to the FSHD phenotype

Our previous MERFISH spatial transcriptomics study revealed heterogenous expression of DUX4 target genes across individual myotubes and identified two possible subpathways of DUX4 target genes represented by *LEUTX* and *ZSCAN4* (Chen et al. 2024). Our current study revealed distinct phenotypes of depletion and overexpression of LEUTX and ZSCAN4, further substantiating the functional differences of DUX4 target subpathways. Interestingly, depletion of KLF18, another DUX4 target gene, also affects expression of a subset of DUX4 target genes, but not *H3.X/Y*, *ZSCAN4* and *LEUTX*, suggesting another subpathway and complexity of DUX4 gene network regulation (Hamm et al. 2026). Coalescing of these distinct DUX4 subpathways may contribute to the observed heterogeneity in FSHD muscle phenotype.

There is a clear apoptosis induction by the DUX4 transgene overexpression (Shadle et al. 2017; Bosnakovski et al. 2018). Neither ZSCAN4 nor LEUTX overexpression in control or FSHD cells, however, exhibits cytotoxicity. This was surprising especially since LEUTX contributes to global upregulation of DUX4-induced genes in FSHD cells. It should be noted that KLF18 depletion was also reported not to rescue DUX4-induced cytotoxicity (Hamm et al. 2026). Consistently, our previous single cell/nucleus transcriptomics analyses of FSHD myotubes failed to show any significant apoptotic gene expression signatures especially later in myotube differentiation when FSHD-induced DUX4 target gene expression is high (Chau et al. 2021; Kong et al. 2024). Together, these results suggest that FSHD-induced DUX4 downstream effectors do not recapitulate the cell death phenotype caused by overexpressed DUX4.

### Dual roles of LEUTX in gene activation and repression to interfere with myotube differentiation/maturation

We recently showed that FSHD myotubes are defective in adaptive response to electrical pulse stimulation (EPS) and that expression of LEUTX alone in control cells is sufficient to recapitulate the phenotype (Kong et al. 2025). Thus, LEUTX-mediated interference of proper myotube differentiation appears to also affect their proper maturation, and has a significant impact on the functionality of FSHD myocytes. Our current results extend this model by demonstrating that LEUTX binds to the promoter and downstream gene body regions of subsets of mitotic/cell cycle and muscle-related genes, and appears to directly regulate them as both an activator and repressor, respectively. Interestingly, LEUTX was found to interact with cohesin in iPSCs (Gawriyski et al. 2023). Cohesin mediates long-distance chromatin interactions, such as topologically associated domain organization and enhancer-promoter interactions, but also binds to the gene body and blocks transcription (Gomes and Espinosa 2010; Chien et al. 2011; Watrin et al. 2016; Chandrasekaran et al. 2022). It is possible that the presence of multiple LEUTX peaks in the downstream gene body regions of repressed targets may also cause unwanted chromatin looping or boundary formation, interfering with RNA polymerase processivity. Further investigation of the dual mechanisms of gene regulation by LEUTX would be important to understand the functional defect of FSHD myocytes.

Because LEUTX normally functions during embryonic development, published binding data from human pluripotent stem cells (hPSCs) (Gawriyski et al. 2023) provide a useful reference for understanding its aberrant activity in FSHD muscle. LEUTX binding pattern differed between PSCs and muscle cells, as well as between LEUTX OE in the control cells and the endogenous LEUTX in FSHD cells. This strongly suggests the context-dependent binding of LEUTX dictated by epigenetic environment and/or cofactor availability. Notably, LEUTX binding peaks in muscle cells, but not in PSCs, were enriched for muscle-specific bHLH transcription factors, such as MYF5, MYOD and MYOG, raising the possibility that these TFs may interact with and function together, or be competed by, LEUTX. Further analyses of binding partners and regulatory mechanism(s) of LEUTX binding would be important to define the mechanisms underlying the context-specific LEUTX activity.

### LEUTX enhances DUX4 target gene expression through DUX4-induced promoter binding

We previously showed that LEUTX depletion and overexpression in FSHD mutant myotubes decreases and enhances global DUX4 target expression, respectively (Kong et al. 2024). However, the underlying mechanism remained unknown. Our CUT&Tag results now demonstrate that LEUTX specifically binds to the DUX4 target gene promoter regions only in FSHD myocytes, but not in control myocytes. LEUTX binding peaks coincide with those of DUX4, strongly suggesting that the initial DUX4 binding enables subsequent LEUTX access, which is missing in control cells. This model is consistent with DUX4 acting as a pioneer factor that recruits p300 coactivator with histone acetyltransferase activity thereby increasing H3 acetylation and promoting open chromatin state (Choi et al. 2016). Notably, a known LEUTX target and DUX4-induced *VMO1* gene promoter is bound by LEUTX in control cells (in the absence of DUX4) coinciding with open chromatin marks (H3K4me3 and H3K27Ac) in control cells, indicative of open chromatin. In contrast, those DUX4 target promoters that are bound by LEUTX only in FSHD cells are lacking these signals in control myocytes, suggesting that the region is closed and inaccessible to LEUTX in the absence of DUX4. Our recent Nanopore long-read genomic sequencing also revealed the general trend of DNA methylation loss in the DUX4 target gene promoters in FSHD mutant cells compared to the isogenic control (Sakr et al. 2025). Together, we propose a model of a dual role of DUX4 in a two-step mechanism of DUX4 target amplification: DUX4 acts as a disease-associated pioneer factor to initiate promoter opening and initial activation of *LEUTX* (and other target genes), which then enables subsequent LEUTX binding and activation (Fig. 7B). We previously showed that histone variants *H3.X/Y* and *LEUTX* stimulate each other’s expression and are both part of the feedforward mechanism of DUX4 target gene network amplification (Kong et al. 2024). Since H3.X/Y maintains the open chromatin state (Resnick et al. 2019), one can envision that the presence of H3.X/Y may also contribute to LEUTX access to DUX4-activated genes.

## Conclusion

Our comparative analyses of two major DUX4 targets, LEUTX and ZSCAN4, representing the two subpathways of DUX4 target gene network revealed their distinct effects on FSHD transcriptomic phenotype. Namely, LEUTX, but not ZSCAN4, amplifies DUX4 target gene activation and interferes with timely cell cycle exit and myotube differentiation by activating mitosis-related genes and repressing muscle-related genes. The results demonstrate the critical role of LEUTX as a major DUX4 effector in FSHD pathogenesis and suggest that unequal expression of functionally distinct DUX4 target genes contributes to disease complexity and phenotypic heterogeneity.

## Limitations of the study

The study focused on LEUTX. Further exploration of ZSCAN4 function and possible combinatorial and individual effects of other target genes are not addressed in the current study, which would be important to understand the disease pathology.

## Methods and materials

### Cell culture, myotube differentiation, and cell viability measurement

Immortalized healthy control 1 (B0, also referred to as CTRL), control 2 (D0), FSHD2 (B2), FSHD1 (B1, C1), and double mutant (DEL4-SM-A, also referred to simply as mutant) were cultured in high glucose DMEM (Gibco) supplemented with 20% FBS (Omega Scientific, Inc.), 1% Pen-Strep (Gibco), and 2% Ultroser G (Crescent Chemical Co.). In the case of the FSHD1 (B1) and the FSHD2 (B2), the identical cell lines were used in (Chau et al. 2021; Kong et al. 2024). Briefly, FSHD1 (B1) carries the 4qA haplotype with 6 D4Z4 repeats, FSHD1 (C1) carries the 4qA haplotype with 2 D4Z4 repeats, and FSHD2 carries a heterozygous *SMCHD1* mutation with two 4qA alleles harboring 15 and 19 D4Z4 repeats, respectively. The double mutant (DEL4-SM-A) carries a homozygous *SMCHD1* mutation and the 4qA haplotype with 2 D4Z4 repeats. Immortalization was performed by following the published protocol in (Shiomi et al. 2011). Briefly, hTERT, p16INK4a-resistant R24C mutant CDK, and cyclin D1 were used to achieve the immortalization. Then, the cells were positively selected for CD56. For stable (cell) clone preparation, cell suspension was distributed in a 96-well plate, and specific clones were selected based on doubling time as well as myotube differentiation efficiency. To differentiate immortalized myoblasts, high-glucose DMEM supplemented with 2% FBS, 1% Pen-Strep (Gibco), and ITS supplement (insulin 0.1%, 0.000067% sodium selenite, 0.055% transferrin; 51300044; Invitrogen) was used where the approximate seeding density of 80 – 90% confluency was used for myotube differentiation. During the myotube differentiation, fresh medium was daily provided for each well. Cell viability was determined by measuring ATP levels using the CellTiter-Glo 2.0 assay (Promega), according to the manufacturer’s instructions.

### Antibodies and primers

The information is in Supplemental Table S5 and Supplemental Table S6.

### RNA, cDNA preparation, and total protein extraction

Total RNA was extracted using RNeasy® Plus Mini (Qiagen) following manufacturer’s protocol. Concentration of extracted RNA was measured by Qubit™ fluorometer (Thermo Fisher Scientific) and Qubit™ RNA High Sensitivity (HS) kit (Thermo Fisher Scientific). cDNA was prepared through reverse transcription using SensiFAST™ cDNA Synthesis Kit (Bioline) following manufacturer’s protocol. cDNA volume was further adjusted with nuclease free water. Total protein was extracted using sample buffer which contains 0.5 % v/v beta-mercaptoethanol in 2X Laemmli Sample Buffer (Bio-Rad). The sample buffer was directly added onto cells and filtered through QIAquick® Spin Columns (Qiagen). The filtered samples were then boiled at 95 °C for 10 minutes.

### RT-qPCR and western blot

qPCR was performed using CFX96 Touch Real-Time PCR Detection System (Bio-Rad) and Azura View Green Fast qPCR Blue Mix LR (Azura Genomics) reagent. Cq values were used to analyze qPCR experimental results, where *GAPDH* Cq values were used to normalize relative gene expression of gene of interest. The Cq values were processed using tidyqpcr package (version 1.0) in R (version 4.3.0), where normalized expression was calculated by 2^(“Δ*Cq*)^. Western blot was performed following a standard protocol. Briefly, 4–20% Mini-PROTEAN® TGX Stain-Free™ Protein Gels (Bio-Rad) was used with Rainbow™ Molecular Weight Markers (Cytiva). Each gel was run at 150 V for 1 hour in 1X running buffer composed of Tris/Glycine/SDS (Bio-Rad). Then, proteins in the gel were transferred to a nitrocellulose membrane, which was later blocked with SuperBlock™ Blocking Buffer (Thermo Fisher Scientific). Then, the membrane was probed with a primary antibody solution then with a secondary antibody conjugated with HRP. For substrate, SuperSignal™ West Pico PLUS Chemiluminescent Substrate (Thermo Fisher Scientific) was used. The membrane was imaged using LAS 4000 (GE Healthcare) for chemiluminescence.

### Plasmid cloning for overexpression experiments

*LEUTX* or *ZSCAN4* gene was cloned into pCW57.1 (Addgene 41393). cDNA of LEUTX including a FLAG tag in overhangs was amplified by PCR using Phusion High-Fidelity PCR Master Mix (Thermo Fisher Scientific). Cloning was achieved using NEBuilder HiFi DNA Assembly Cloning Kit (New England Biolabs), following the manufacturer’s protocol. The FLAG tag was added to the C-terminus of LEUTX cDNA and to the N-terminus of ZSCAN4 cDNA, avoiding DNA binding site of each protein. EcoRI and SalI were used as restriction enzymes (New England Biolabs). cDNA fragments as well as plasmids were sequenced to confirm the cloning.

### Lentivirus packaging for depletion and overexpression experiments

For depletion experiments, shRNA plasmids (MISSION shRNA, Sigma-Aldrich) were used and packaged into lentivirus. ShRNA sequences are listed in Table S6. For the overexpression experiments, pCW57.1 (Addgene 41393) was used for cloning and lentivirus packaging. To package lentivirus, lentiviral transfer plasmid, PAX2 (Addgene #12260), and VSVG (Addgene #8454) were co-transfected into 293T cells at a ratio of 4:3:2 using Lipofectamine 3000 (Thermo Fisher Scientific). The medium was changed 8 hours after the co-transfection. Lentivirus was harvested twice, one after 24 hours and another after 48 hours of co-transfection. Collected lentiviruses were aliquoted, snap-frozen in liquid nitrogen, and stored in -80 °C. The same method was used for preparing lentivirus delivering Tet-inducible gene of interest for overexpression. In the case of lentivirus infection for depletion experiments, myoblasts were infected with the lentivirus for 8 or 24 hours. After the infection, the viral medium was replaced with the growth media. Puromycin, at the final concentration of 3 μg/mL, was used as a selection marker and therefore to confirm the lentivirus infection. In the case of lentivirus infection for overexpression experiments, either a single clone of stable cell line or transiently selected heterogenous population by puromycin was used. DOX was used to induce LEUTX overexpression. In LEUTX OE RNA-seq and CUT&Tag experiments, 1 μg/mL DOX was added 48 hours prior to harvest at differentiation day 4. The concentration and timing for other experiments are indicated in the figures or figure legend.

### 5-ethynyl-2’-deoxyuridine (EdU) labeling and detection

To detect cells in S phase, EdU was pulse labeled in cell culture by using Click-iT™ EdU Cell Proliferation Kit for Imaging, Alexa Fluor™ 488 dye reagent (Thermo Fisher Scientific) following the manufacturer’s protocol. Briefly, samples were treated with 10 μM EdU for 3 hours before fixing with 4 % v/v paraformaldehyde. The EdU signals were detected by fluorescence microscopy.

### Immunofluorescent staining

To prepare samples for fluorescence imaging, cells were cultured either on 12 mm diameter cover glasses or either 4-well or 8-well Lab-Tek chamber slides (Thermo Fisher Scientific). Both cover glasses and chamber slides were pre-treated with poly-L-lysine to coat the surface before seeding cells. To stain the samples with antibody, a standard immunofluorescence protocol was used. Briefly, cells were fixed with 4 % v/v paraformaldehyde for 10 minutes at room temperature, permeabilized with 0.5 % v/v Triton X-100 in 1X PBS for 10 minutes at 37 °C., and blocked for 15 minutes at 37 °C. To prepare both blocking and antibody solution, a base solution called SNBP (0.02 % v/v saponin, 0.05 % v/v sodium azide, and 1 % m/v BSA) was prepared first. The blocking solution was prepared by adding 4 % v/v horse serum and 0.1 % v/v gelatin in SNBP. The antibody solution was prepared by adding 1 % v/v horse serum and 0.05 % gelatin in SNBP, which was used as a diluent of stock antibodies. After blocking, samples were treated with either a primary or a cocktail of primary antibodies at 4 °C overnight. Then, samples were washed three times with 0.005 % v/v Triton X-100 in 1X PBS (PBST). Then, samples were treated with either a secondary or a cocktail of secondary antibodies at 37 °C for 30 minutes. Finally, samples were washed twice and stained with DAPI for 3 minutes. Fluoromount-G™ Mounting Medium (Thermo Fisher Scientific) was used to apply seals on the sample. For image acquisition, Keyence BZ-X800 fluorescence microscope was used.

### Fluorescence imaging analysis

Fluorescence images were both manually and automatically analyzed by ImageJ (FIJI) or publicly available software. To judge and determine positive signals of LEUTX and H3.X/Y, the nuclei clusters were manually selected first and then intensity of LEUTX and H3.X/Y signals were evaluated by two researchers for cross checking. In the case of fusion index and subsequent myotube morphology related information, 10X fluorescence images and MyoFInDer software (Weisrock et al. 2024) was used. For DAPI and EdU counting, 10X fluorescence images were used. To analyze DAPI and EdU signals, custom ImageJ macro scripts were prepared. Briefly, for DAPI channel, the threshold was set from 20 to 255, and for EdU channel, the threshold was set from 10 to 255. To avoid nuclei clusters when analyzing EdU positive signals from unfused myoblasts, a circularity threshold of 0.6 – 1.0 was used in DAPI channel for region of interest (ROI) registration. For both channels, any objects whose area is under 0.001 were excluded. In the case of the LEUTX and KI67 fluorescence intensity measurement, ROIs within top 25% mean fluorescence intensity were selected for analysis.

### Bulk RNA-seq library preparation and analysis

Total RNA was extracted using RNeasy® Plus Mini (Qiagen) following manufacturer’s protocol. To prepare full length cDNA for the library preparation, the Smart-Seq2 protocol with minor modifications was followed (Picelli et al. 2014; Jiang et al. 2020). The library was then prepared using Nextera DNA Flex (Illumina) kit. The sequencing was performed on the Illumina NextSeq500 platform. For pseudoalignment and quantification, Kallisto (Bray et al. 2016) (version 0.48.0) was used where human genome reference hg38 cDNA was used as a reference. For differential gene expression (DEG) analysis, Sleuth (Pimentel et al. 2017) (version 0.30.1) was used. Briefly, the gene mode was used in the Sleuth software, and the beta values were used as fold-changes values after transforming into log_2_ scale. +1 and -1 were used as threshold values for the fold-change in log_2_ scale. q-value threshold value was set to 0.1 for all DEG analyses. For GO term analysis, ClusterProfiler (Yu et al. 2012) (version 4.6.2) was used. Visualization of bulk RNA-seq data was achieved by ggplot2 (version 3.5.1.9000), ggpubr (version 0.6.0), and ComplexHeatmap (Gu et al. 2016) (version 2.14.0) packages. For the motif analysis, HOMER (version 4.11) was used. R (version 4.2.2 or version 4.3.0) was used.

### CUT&Tag

CUT&Tag assays for LEUTX and H3K4me3 were performed using the CUT&Tag Assay Kit (Cat. No. 77552, Cell Signaling Technology, MA, USA) according to the manufacturer’s instructions. Control 1 (B0) cells harboring a stable doxycycline-inducible LEUTX expression system or FSHD1 (C1) cell line with high DUX4 target gene expression were seeded at 5 × 10⁵ cells per well in 6-well plates. Cells were differentiated the following day and harvested on differentiation day 4 after 48 hours of doxycycline induction. For each assay, 10% of the harvested cells were lysed, and the lysate was incubated with Con A–coated magnetic beads at room temperature for 1 hour. Samples were then sequentially incubated at room temperature with primary antibody or pre-immune IgG for 1 hour, followed by a 30-minute incubation with a secondary antibody and a 1-hour incubation with pA/G-Tn5 for binding. Tagmentation was performed at 37 °C for 1 hour in tagmentation buffer. Tagmented chromatin fragments were released, purified, and used for library preparation. Indexed libraries were pooled and sequenced on a NovaSeq X Plus platform (Illumina, CA, USA) with paired-end 150 bp reads. Each LEUTX antibody condition was performed with two biological replicates. Adapter trimming and quality filtering were performed using fastp (v0.23.2). High-quality clean reads were aligned to the hg38 reference genome using Bowtie2 (v2.5.1). Peak calling was performed using MACS2 (v2.2.7.1) with matched IgG controls. Peaks with a q value < 0.01 and a fold enrichment ≥ 3.0 were considered significant. High-confidence peaks were defined as those present in both biological replicates and were used for downstream analyses. Peak annotation was performed using ChIPSeeker (v1.42.1), and motif enrichment analysis was carried out using HOMER (v5.1).

## Statistical analysis

Unless otherwise specified, all statistical tests were performed using rstatix (version 0.7.2) packages in R (version 4.2.2 or 4.30). Specific statistical tests and information are noted in the figure legends.

## Data availability

The RNA-seq data and CUT&Tag data generated in this study are available in The database of Genotypes and Phenotypes (dbGaP; https://www.ncbi.nlm.nih.gov/gap/) with accession number phs002554 (The submission to dbGaP is still in progress). The DUX4 ChIP-seq dataset (GEO accession number GSE33838; Geng et al., 2012) was also used in this study.

## Acknowledgements

We are very grateful to Sean R. Bennett, Danielle C. Hamm and Stephen Tapscott at Human Biology Division, Fred Hutchinson Cancer Center for sharing the monoclonal antibody specific for LEUTX. We appreciate Heidi Liang for operating the sequencer and processing the libraries. We also appreciate Jasmine Sakr for insights and support on sequencing data analysis and thank Elnaz Abdollahzadeh for manuscript editing. This work was funded by National Institutes of Health grant R01AR071287 (KY and AM).

---

**Supplemental Figure 1.**
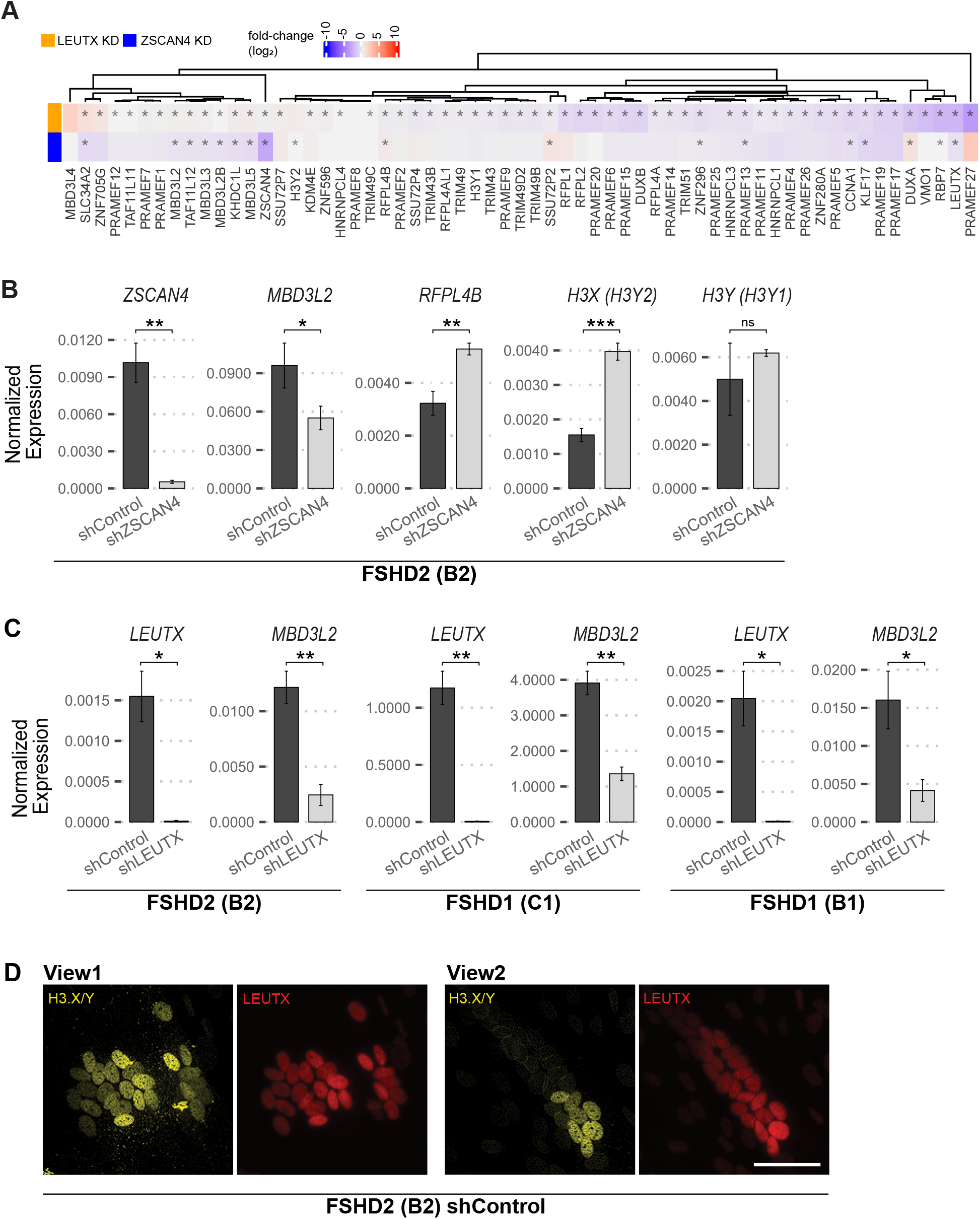
Bulk RNA-seq and RT-qPCR results of DUX4 target genes from LEUTX and ZSCAN4 depletion experiments and representative immunofluorescence images of LEUTX and H3.X/Y colocalization. **A.** A heatmap of fold-change values obtained from the bulk RNA-seq experiments between the LEUTX depletion and the ZSCAN4 depletion in the mutant myotubes is shown. The heatmap shows 61 pre-selected DUX4 targets where asterisks indicate statistically significant (q-value < 0.1) fold-change values. **B.** RT-qPCR results of ZSCAN4 depletion in the FSHD2 (B2) are shown. ZSCAN4, MBD3L2, RFPL4B, H3.X, and H3.Y expression levels were measured. GAPDH was used to normalize the expression level. n=3. Student’s t-test. * p ≤ 0.05, ** p ≤ 0.01, *** p ≤ 0.001, and ns not significant. **C.** RT-qPCR results of LEUTX depletion in the FSHD2 (B2), the FSHD1 (C1) and the FSHD1 (B1) are shown. MBD3L2 was used as a proxy for the DUX4 target genes. GAPDH was used to normalize the expression level. n=3. Student’s t-test. * p ≤ 0.05 and ** p ≤ 0.01. **D.** Digitally zoomed in 20X immunofluorescence images of H3.XY and LEUTX positive signals obtained from the FSHD2 (B2) myotubes are shown. Scale bar = 50 μm.

**Supplemental Figure 2.**
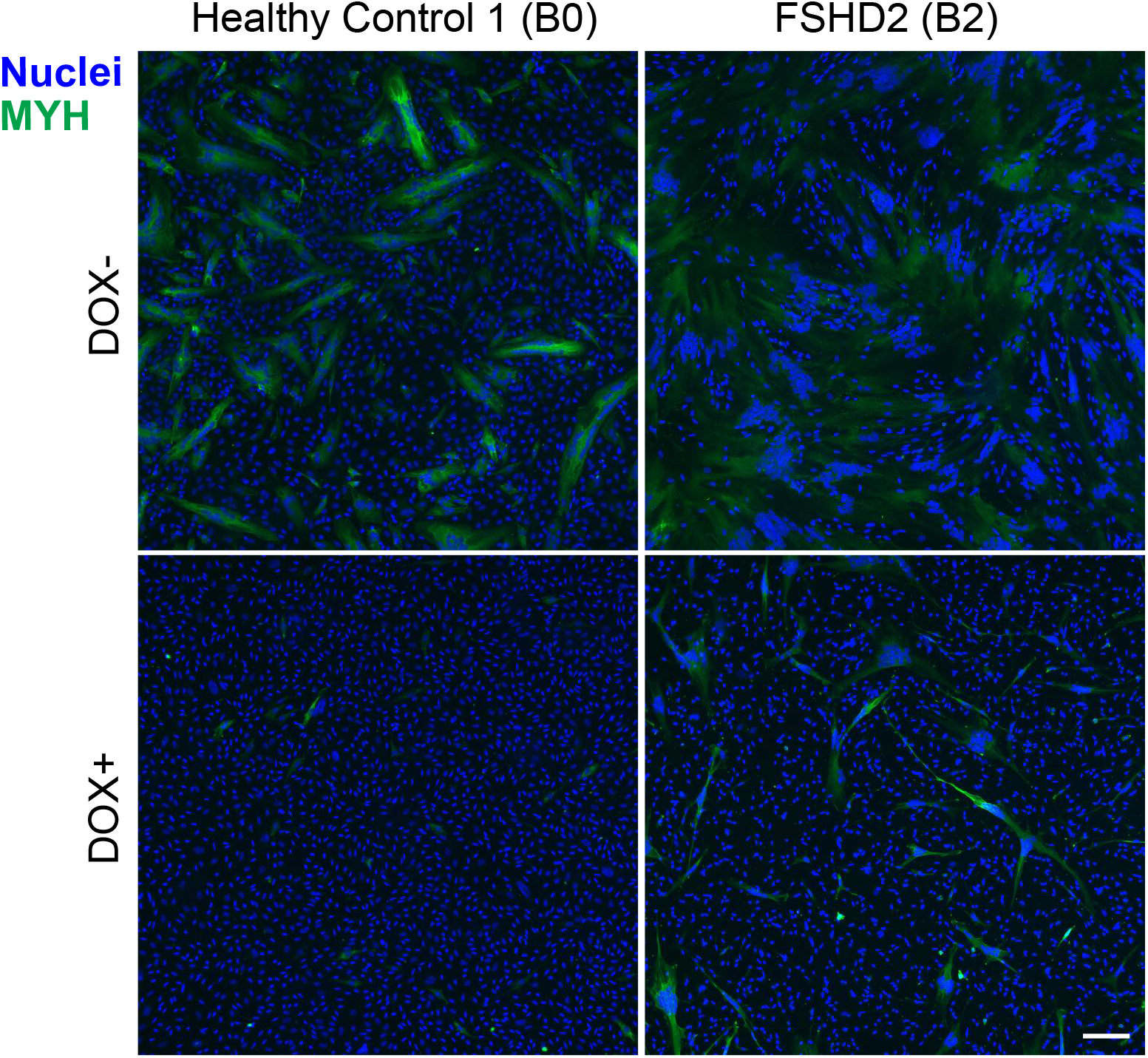
Representative Immunofluorescence images to indicate inhibited myotube differentiation with LEUTX OE. MYH staining indicates the location of myotubes. DAPI was used to stain nuclei. Representative images with and without DOX treatment from the two cell lines are shown here. Scale bar 100 μm.

**Supplemental Figure 3.**
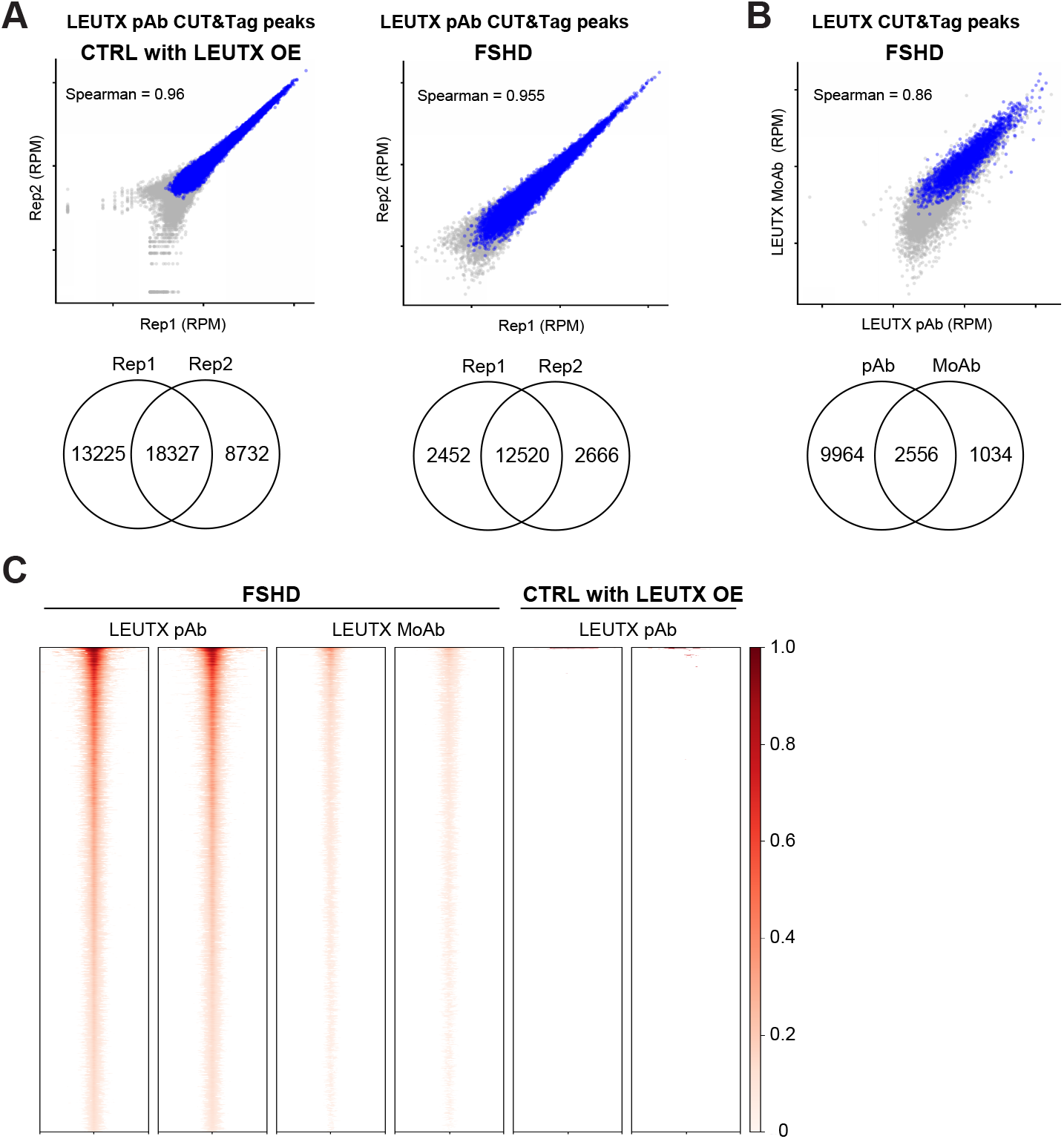
LEUTX monoclonal antibody confirms the specificity of the polyclonal LEUTX antibody binding sites in FSHD cells. **A**. Scatter plots showing LEUTX CUT&Tag peak counts (RPM) in two independent replicates of DOX-inducible LEUTX Control myocytes (left) and FSHD myocytes (right). Each point represents an individual LEUTX peak, with blue dots indicating common peaks between the two replicates. Corresponding Venn diagrams show the overlap of LEUTX-binding sites between the two replicates. **B**. Same as A. Scatter plots showing the correlation of LEUTX peak counts at common peaks identified in FSHD myocytes using monoclonal and polyclonal anti-LEUTX antibodies. The corresponding Venn diagrams are also shown. **C**. Heatmap comparison of LEUTX binding signals. LEUTX pAb peaks in FSHD enLEUTX myocytes were ranked from strongest to weakest and compared with LEUTX MoAb signals in FSHD enLEUTX myocytes and LEUTX pAb signals in Control LEUTX OE myocytes at the corresponding regions. Relative signal densities, scaled from 0 to 1 as indicated by the color bar, are shown within a 4-kb window centered on each LEUTX peak. Two independent replicates are shown for each condition.

**Supplemental Figure 4.**
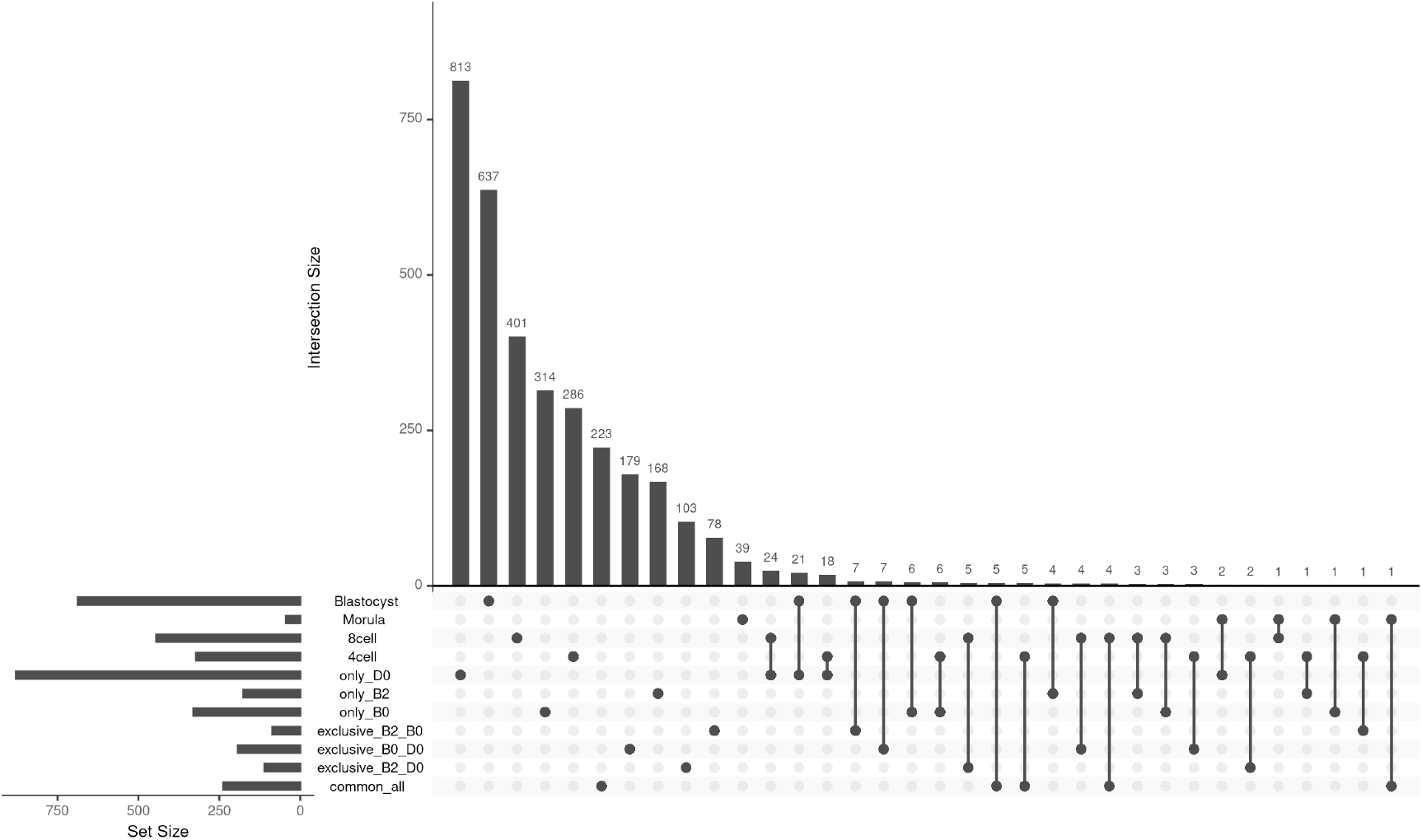
Upregulated DEGs by LEUTX OE have minimal overlap with ZGA signature genes (Yan et al. 2013; Zheng et al. 2024). Each dot represents an individual gene set, and connected dots represent specific intersections among sets. The top bar graph shows the size of each intersection, while the bottom bar graph shows the size of each individual set.To generate the sets used in this upset plot, a three-way Venn diagram analysis was first performed using upregulated DEGs from LEUTX OE across three cell lines (B0 control, D0 control, and B2 FSHD2). The upset plot then displays the number of overlapping genes between these LEUTX-induced upregulated DEGs and ZGA signature gene sets (4-cell, 8-cell, Morula, and Blastocyst).

**Supplemental Figure 5.**
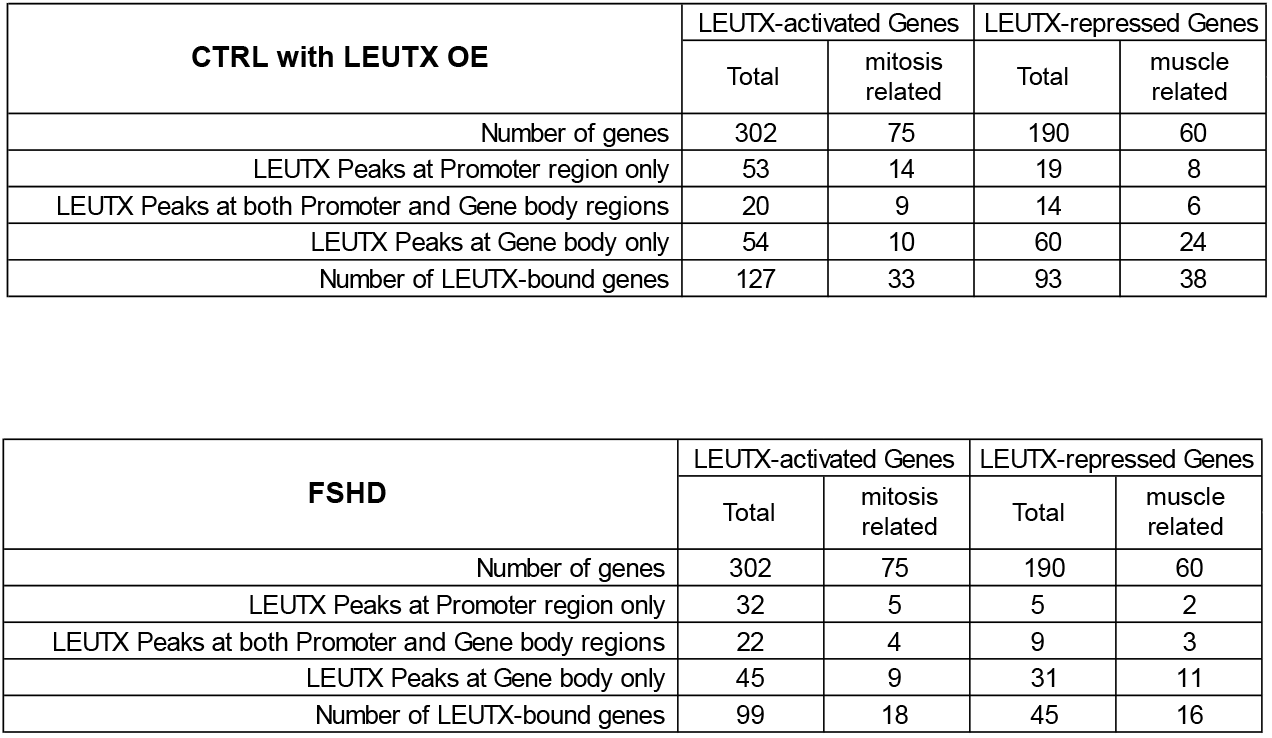
Distribution of LEUTX binding peaks across gene regions in common differentially expressed genes (DEGs) upon LEUTX knockdown. Summary of LEUTX binding peak distribution in Control with LEUTX OE and FSHD cells across different gene regions of common DEGs identified in LEUTX knockdown FSHD2 and mutant cells. Genes were categorized based on LEUTX peak localization as promoter only, both promoter and gene body, or gene body only. The number of LEUTX-bound genes in each category is indicated.

**Supplemental Figure 6.**
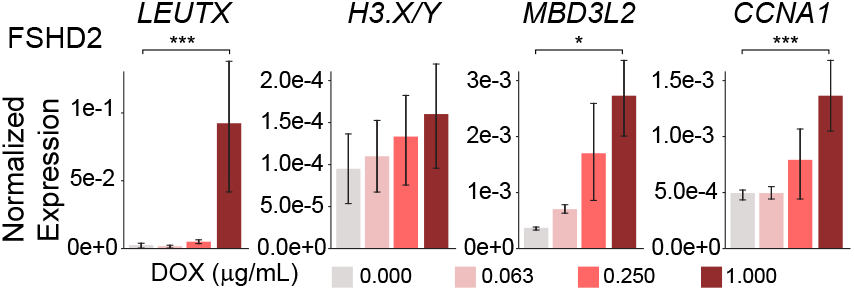
LEUTX overexpression enhanced DUX4 target gene expression in patient cells. LEUTX overexpression was modulated by DOX concentration titration in the FSHD2 myotubes. RT-qPCR was performed to measure the expression level of DUX4 targets (LEUTX, H3.X/Y, MBD3L2, and CCNA1). 1X DOX indicates the final concentration of 1 μg/mL. Student’s t-test. * p ≤ 0.05, *** p ≤ 0.001. n=4.

